# A genetic screen identifies a role for *oprF* in *Pseudomonas aeruginosa* biofilm stimulation by subinhibitory antibiotics

**DOI:** 10.1101/2023.09.05.556382

**Authors:** Luke N. Yaeger, Michael R. M. Ranieri, Jessica Chee, Sawyer Karabelas-Pittman, Madeleine Rudolph, Alessio M. Giovannoni, Hanjeong Harvey, Lori L. Burrows

## Abstract

Biofilms are surface-associated communities of bacteria that grow in a self-produced matrix of polysaccharides, proteins, and extracellular DNA (eDNA). Sub-minimal inhibitory concentrations (sub-MIC) of antibiotics induce biofilm formation, indicating a potential defensive response to antibiotic stress. However, the mechanisms behind sub-MIC antibiotic-induced biofilm formation are unclear. We show that treatment of *Pseudomonas aeruginosa* with multiple classes of sub-MIC antibiotics with distinct targets induces biofilm formation. Further, addition of exogenous eDNA or cell lysate failed to increase biofilm formation to the same extent as antibiotics, suggesting that the release of cellular contents by antibiotic-driven bacteriolysis is insufficient. Using a genetic screen to find stimulation-deficient mutants, we identified the outer membrane porin OprF and the extracytoplasmic function sigma factor SigX as important for the phenotype. Similarly, loss of OmpA – the *Escherichia coli* OprF homologue – prevented sub-MIC antibiotic stimulation of *E. coli* biofilms. The C-terminal PG-binding domain of OprF was dispensable for biofilm stimulation. Our screen also identified the periplasmic disulfide bond-forming enzyme DsbA and a predicted cyclic-di-GMP phosphodiesterase encoded by PA2200 as essential for biofilm stimulation. The phosphodiesterase activity of PA2200 is likely controlled by a disulfide bond in its regulatory domain, and folding of OprF is influenced by disulfide bond formation, connecting the mutant phenotypes. Addition of the reducing agent dithiothreitol prevented sub-MIC antibiotic biofilm stimulation. Finally, we show that activation of a c-di-GMP-responsive promoter follows treatment with sub-MIC antibiotics in the wild-type but not an *oprF* mutant. Together, these results show that antibiotic-induced biofilm formation is likely driven by a signalling pathway that translates changes in periplasmic redox state into elevated biofilm formation through increases in c-di-GMP.

**Significance:** Bacterial biofilms cause significant treatment challenges in clinical settings due to their ability to resist antibiotic killing. Paradoxically, sub-inhibitory levels of a number of chemically distinct antibiotics, with different modes of action, can stimulate biofilm formation of multiple bacterial species. In this work we used the biofilm forming opportunistic pathogen *Pseudomonas aeruginosa* to look for mutants blind to sub-inhibitory antibiotic exposure, failing to demonstrate biofilm stimulation in response to 3 different antibiotics. We identified key players in this response, including the outer membrane porin OprF and the sigma factor SigX. Notably, the hits from our mutant screen connect changes in periplasmic redox state to elevated biofilm formation via c-di-GMP signaling. By understanding these underlying mechanisms, we can better strategize interventions against biofilm-mediated antibiotic resistance in *P. aeruginosa* and other bacterial species.

## Introduction

*Pseudomonas aeruginosa* is a Gram-negative, opportunistic pathogen and a leading cause of nosocomial infections. The World Health Organization identified it as a critical priority pathogen due to its multi-drug resistance and ability to form biofilms.^1–3^ Patients with cystic fibrosis are especially susceptible to chronic *P. aeruginosa* infections; mutations in the cystic fibrosis transmembrane conductance reporter (*CFTR*) gene produce an environment conducive to infection with *P. aeruginosa*.^4^ Biofilm formation by *P. aeruginosa* is a major factor in colonization of various surfaces including medical devices and subsequent recalcitrance to treatment, resulting in chronic infections that are very difficult to eradicate. The identification of new therapies to combat *P. aeruginosa* infections is paramount for successful treatment, and understanding biofilm biology can help guide more effective therapeutic design.

Biofilms are surface-associated microbial communities that grow in a matrix of self-produced extracellular polymeric substances (EPS). The EPS contains polysaccharides, lipids, proteins, and extracellular DNA (eDNA). There are three secreted polysaccharides found in *P. aeruginosa* biofilms.^5^ Psl is a repeating pentasaccharide of (3x) D-mannose, D-glucose, and L-rhamnose whose biosynthetic machinery is encoded by the *pslA-O* operon.^6^ Pel is synthesized by products of the operon *pelA-F* and is a repeat of partially acetylated, 1-4 linked N-acetylgalactosamine and N-acetylglucosamine.^7,8^ Both Psl and Pel play roles in early biofilm formation by facilitating attachment to a surface, as well as acting as structural components in mature biofilms.^8–11^ Alginate is an anionic polysaccharide with strain- and condition-dependent expression in biofilms;^12^ its production is induced in the host during chronic infection by exposure to reactive oxygen species produced by immune cells.^13^ Alginate production contributes to a mucoid phenotype associated with increased antibiotic tolerance and immune evasion.^14–16^

*Pseudomonas* biofilm formation begins upon association with a surface, which is termed “reversible attachment”. Reversible attachment is mediated by flagella and pili at the cell pole and causes perpendicular association of cells with the surface.^17^ When forming a biofilm, levels of the secondary messenger cyclic di-GMP (c-di-GMP) increase and the cell irreversibly attaches to the surface by its longitudinal axis via Psl or Pel polysaccharides acting as adhesins.^11,17^ Growth on the surface occurs along with the production of EPS to form microcolonies that later become a mature biofilm. Mature biofilms are highly ordered, and interactions between eDNA and Pel polysaccharide provide structural integrity.^8,18,19^

Biofilms confer many survival advantages, including the sharing of resources, environmental protection, antibiotic tolerance, and antibiotic resistance.^20–22^ Nutrient, oxygen, and pH gradients form within a biofilm, resulting in phenotypic heterogeneity.^21,23^ Cells that grow more slowly in some areas of those gradients are tolerant to antibiotic treatment, leading to relapse of infection. The proximity and reduced motility of cells in biofilms allows for increased horizontal gene transfer, which may increase spread of resistance genes.^22^ The hydrated matrix prevents desiccation and creates a barrier that can slow the passage of certain toxic molecules, including antibiotics.^24,25^ While the properties of biofilms are well characterized, the diverse cues that modulate biofilm formation continue to be discovered. For example, exposure to sub-lethal antibiotics is potent driver of *P. aeruginosa* biofilm formation,^26^ but the pathway from antibiotic treatment to biofilm formation is poorly understood.

The ability of antibiotics to act as signaling molecules has been recognized.^27–29^ They are produced in the environment at much lower concentrations than those used therapeutically, and it is likely that below their minimum inhibitory concentrations (sub-MIC), these molecules shape single-cell and community behaviours. Global gene expression profiles of multiple organisms are altered in the presence of sub-MIC antibiotics, including genes not directly related to antibiotic mechanism of action or stress/damage pathways.^27,28,30–32^ Examples include tetracycline-dependent induction of cytotoxicity in *P. aeruginosa* via the type III secretion system (T3SS), or azithromycin-dependent downregulation of multiple quorum sensing genes.^27,33^ Sub-MIC antibiotics also influence biofilm formation. For example, sub-MIC levels of tobramycin stimulate biofilm formation by multiple isolates of *P. aeruginosa*.^34,35^ Other antibiotics such as ciprofloxacin and tetracycline – with separate targets and modes of action – have similar effects.^27^ Increased alginate production was reported following treatment with imipenem, norfloxacin, ofloxacin, and ceftazidime.^36,37^ Similar responses to sub-MIC antibiotic exposure have been reported for multiple bacterial species and antibiotic classes, suggesting that the increased biofilm formation may represent a generic defensive response, coordinated through existing stress pathways and/or potentially novel mechanisms.^26^

Although the exact mechanisms underlying antibiotic-induced increases in biofilm formation of *P. aeruginosa* and other bacteria are currently unknown, there are two emerging, non-mutually exclusive hypotheses. The first, known as the seeding hypothesis, posits that sub-MIC antibiotics cause cell death in a susceptible subpopulation of bacterial cells, which leads to release of eDNA and other cell contents, promoting biofilm formation by the remaining cells.^38–41^ The second hypothesis proposes that the increase in biofilm formation is part of a coordinated response to the presence of sub-MIC antibiotics.^27^ This involves the detection of antibiotics, either directly or, more likely, the cellular stresses caused by their action, with an increase in production of adhesins, matrix polysaccharides, and eDNA. There are a few examples that support the concept of a coordinated biofilm stimulation response. For example, cell-surface interactions increase as a result of sub-MIC antibiotic treatment of both *P. aeruginosa* and *S. aureus*, via up-regulation of adhesion-related proteins and changes in cell surface hydrophobicity.^42,43^ Tobramycin induces biofilm formation in a subset of *P. aeruginosa* strains via the c-di-GMP phosphodiesterase Arr;^34,35^ while another investigation of biofilm stimulation by sub-MIC tobramycin showed that eDNA, the *Pseudomonas* Quinolone Signal (PQS), and the iron starvation response small regulatory RNAs *prrF1/F2* were involved.^44^

Despite the identification of some components, there has been no systematic, untargeted screen for factors modulating the *P. aeruginosa* biofilm stimulation response to sub-MIC antibiotic exposure. The conservation of the response to chemically- and mechanistically-distinct antibiotics, in a variety of bacterial species, implies a generalized reaction to the common effects of antibiotic action. In this study, we hypothesized that increased biofilm formation in response to sub-MIC antibiotic exposure is part of a conserved and coordinated stress response, and sought to identify common components required for biofilm stimulation in response to multiple antibiotics.

## Results

### Sub-MIC cefixime, thiostrepton, and tobramycin stimulate biofilm formation

To identify genes required for increased biofilm formation following exposure to antibiotics, we developed a screen for mutants unable to mount that response. First, we optimized the concentration range of three chemically- and functionally-diverse antibiotics – cefixime, tobramycin, and thiostrepton – that stimulate biofilm formation of *P. aeruginosa* strain PAO1, for use in subsequent mutant screening experiments. Cefixime is a cephalosporin that inhibits peptidoglycan cross-linking, tobramycin is an aminoglycoside that disrupts translation, and thiostrepton is a thiopeptide antibiotic that inhibits translation in a manner distinct from tobramycin. Both cefixime and tobramycin are used to treat *P. aeruginosa* infections. Thiostrepton is not used in humans; however, it was selected based on previous work that identified it as a potent stimulator of *P. aeruginosa* biofilm formation.^45^ Based on our past work, we set an arbitrary cut-off of 200% of the vehicle control to define biofilm stimulation. Sub-MIC cefixime, tobramycin, and thiostrepton all stimulated biofilm formation compared to an untreated control (**Figure 1**). For cefixime and tobramycin, the maximal stimulatory concentrations were approximately ¼ to ½ of the MIC. The maximal stimulatory concentration of thiostrepton tested (10 µM) was constrained by its limited solubility.

**Figure 1:**
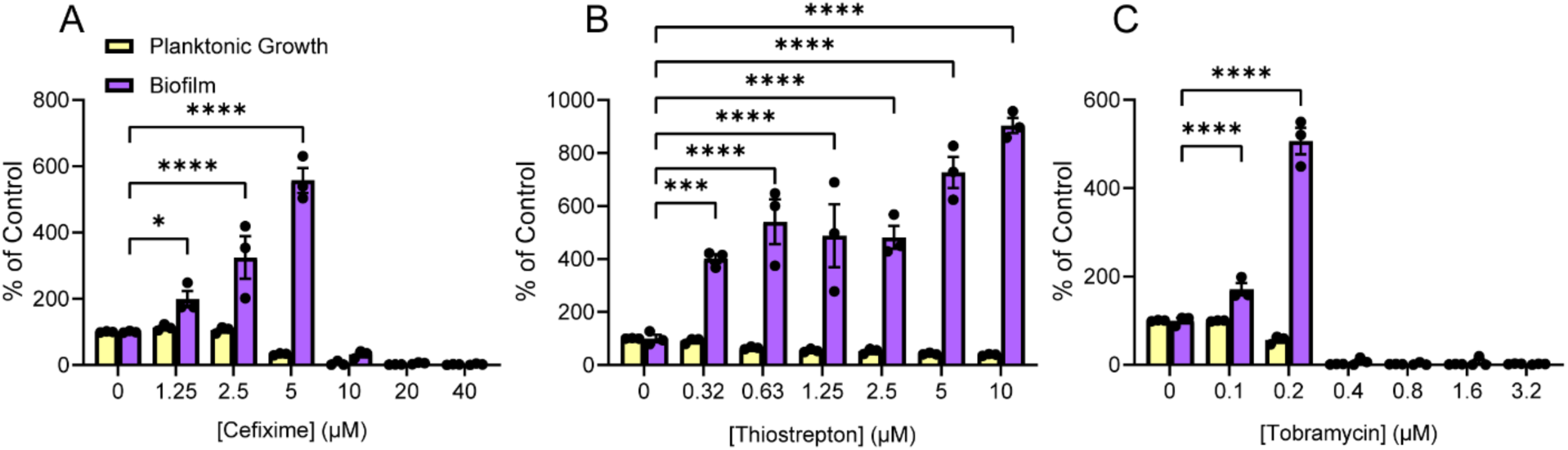
Sub-MIC antibiotics stimulate biofilm formation of *P. aeruginosa* PAO1. Structurally and functionally diverse antibiotics (A) cefixime, (B) thiostrepton, and (C) tobramycin cause dose-dependent increases in biofilm formation as they approach the minimal inhibitory concentration. A two-way ANOVA followed by a Dunnett’s test was performed to compare the untreated control and each drug treatment. * = p<0.05, ** = p<0.01, *** = p<0.001, **** = p<0.0001. Three biological replicates were performed, with 3 technical replicates each. A representative biological replicate is shown, with individual data points shown as black circles and error bars representing the standard error of the mean. Planktonic growth (OD_600_, yellow) and biofilm (A_600_, purple) are both reported as percentage of the vehicle control.

### Neither eDNA nor cell lysate increases biofilm to the same extent as sub-MIC antibiotics

Cefixime and tobramycin are bactericidal antibiotics, while thiostrepton is bacteriostatic. However, thiostrepton caused the largest increase in biofilm formation relative to the DMSO control. This result suggests that cell lysis and release of intracellular contents are less likely to the main drivers of biofilm formation, arguing against the seeding hypothesis. We tested the seeding hypothesis directly by adding purified eDNA to PAO1 cultures, then measuring the amount of biofilm produced compared to control. The eDNA concentration range was based on the amount of DNA per cell combined with estimates of the number of cells undergoing lysis.^46,47^ Specifically, the mass of a single *P. aeruginosa* genome is ∼6.8 x 10^-6^ ng (assuming a 6.3 million bp genome and an average 650 g/mol per base pair). In our assay conditions, there are approximately 1.5 x 10^7^ CFU per mL. Assuming a maximal cell lysis of 50% at ½ MIC, we calculated that the total eDNA released would be ∼51 ng, corresponding to 0.34 ng/µL. Therefore, we selected a concentration range of *P. aeruginosa* genomic DNA that would encompass this estimate. The addition of eDNA in this range did not significantly increase biofilm formation (**Figure 2a**).

**Figure 2:**
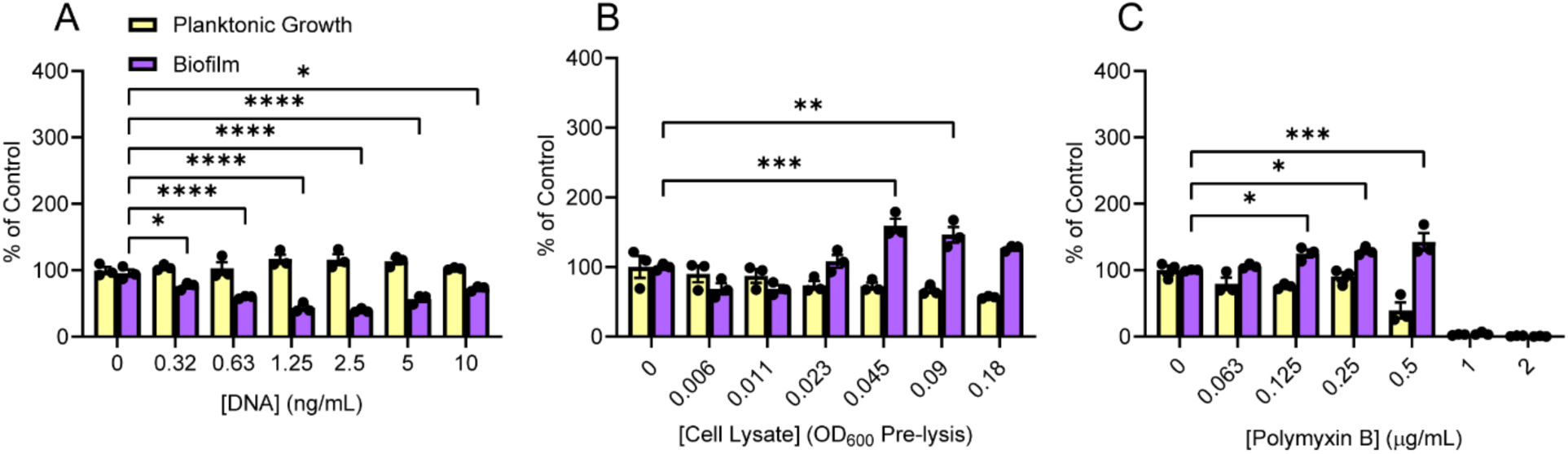
Release of cellular contents fails to recapitulate antibiotic-induced biofilm formation. A) Addition of purified genomic DNA from *P. aeruginosa* PAO1 does not increase biofilm formation at any of the concentrations tested. B) Addition of cell lysate does not induce biofilm formation above our cutoff of 200% of control at any of the concentrations tested, although a small yet significant biofilm induction occurs at higher concentrations of lysate. C) Sub-MIC polymyxin B is a poor inducer of biofilm formation and fails to induce biofilm above 200% of the untreated control, despite a statistically significant increase in biofilm. A two-way ANOVA followed by Dunnett’s test was performed to compare the untreated control and each drug treatment. * = p<0.05, ** = p<0.01, *** = p<0.001, **** = p<0.0001. Three biological replicates were performed with 3 technical replicates each. A representative biological replicate is shown, with individual data points shown as black circles and error bars representing the standard error of the mean. Planktonic growth (OD_600_, yellow) and biofilm (A_600_, purple) are reported as percentage of the untreated-treated control.

In contrast to DNA released by antibiotic-induced lysis, purified genomic DNA might lack associated DNA-binding proteins or other factors released during lysis that stimulate biofilm formation. For example, an unknown factor released upon *P. aeruginosa* lysis was reported to act as a warning signal for kin cells, activating the Gac/Rsm pathway involved in up-regulating type-VI secretion and biofilm formation.^48^ Therefore, we also measured changes in biofilm formation in response to addition of PAO1 whole-cell lysate. Concentrations were based on cell densities at the time of inoculation (∼1.1×10^4^ cfu/mL) and assay endpoint (∼1.9×10^7^ cfu/mL), aiming to include concentrations of cell lysate that could represent 50-75% of cells in the assay undergoing lysis. Addition of whole cell lysate also failed to substantially increase biofilm formation (**Figure 2b**). We observed a small increase in biofilm at the highest concentrations of added lysate, however, the change was minor compared to that resulting from exposure to sub-MIC antibiotics. Finally, addition of subinhibitory concentrations of polymyxin B, a membrane-targeting antibiotic that causes rapid cell lysis, failed to stimulated biofilm formation to the same degree as the other antibiotics tested (**Figure 2c**). Together, these results indicate that release of eDNA and/or cellular contents alone is unlikely to explain the biofilm stimulation observed following treatment with sub-MIC antibiotics.

### Identification of genes involved in the biofilm response to sub-MIC antibiotics

Since the addition of eDNA or cell lysate failed to recapitulate antibiotic-induced biofilm formation, we hypothesized that exposure to sub-MIC antibiotics may activate stress responses that increase biofilm formation. To identify genes involved in such responses, we screened for mutants unable to react to sub-inhibitory concentrations of three different antibiotics with increased biofilm formation (**Figure 3a**). We generated a Mariner transposon library containing approximately 13,000 mutants and, as a first pass, screened for mutants deficient in the biofilm response to ½ MIC of cefixime. Mutants that fell below the 200% of vehicle control cut-off were picked for follow up dose-response biofilm assays using cefixime, tobramycin, and thiostrepton, and those with impaired responses to all three antibiotics were chosen for additional investigation. The site of transposon insertion in each mutant was identified using arbitrarily primed touchdown PCR and sequencing.

**Figure 3:**
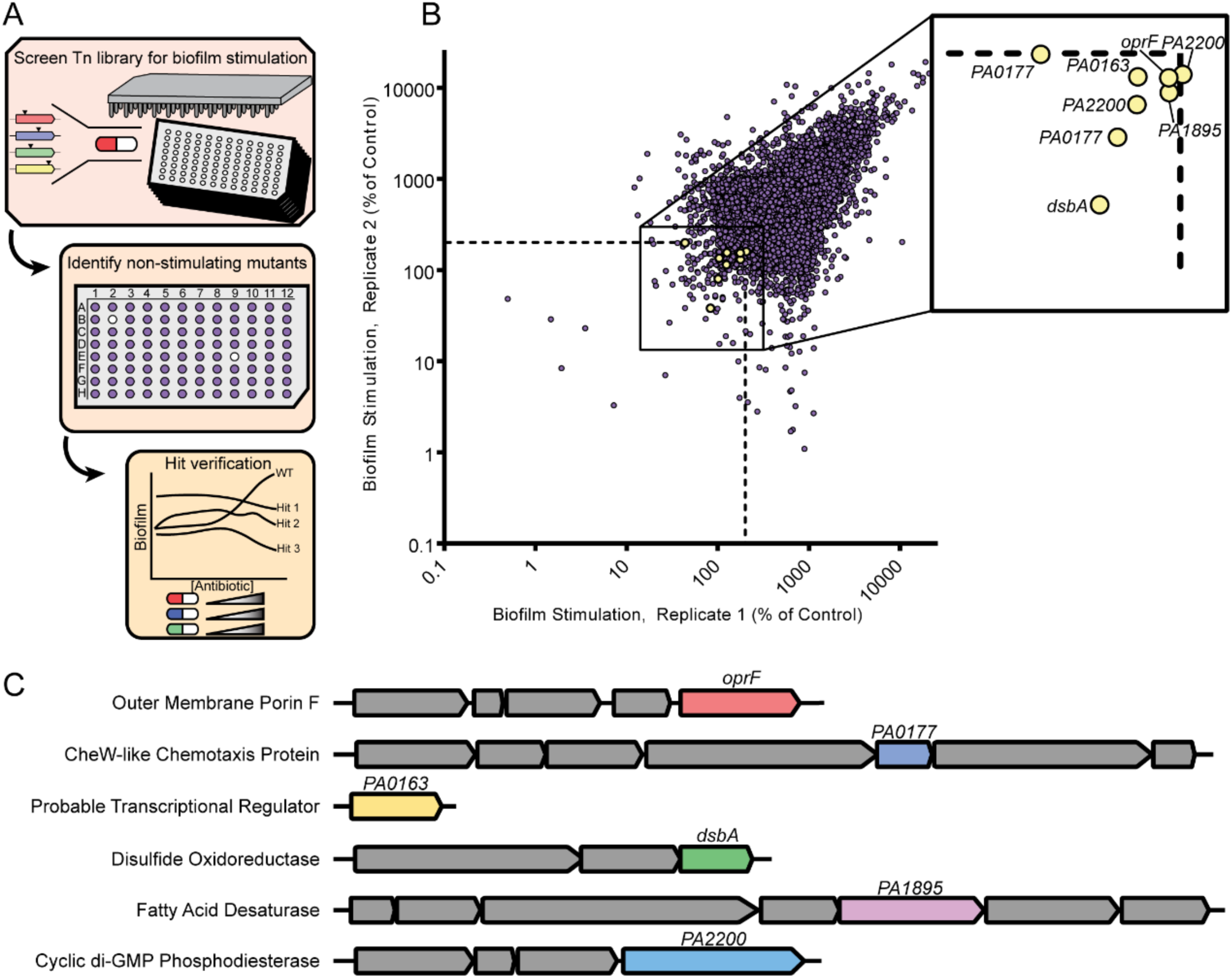
A transposon mutant library screen uncovers genetic determinants of biofilm stimulation by sub-MIC antibiotics. A) A PAO1 transposon library was constructed and arrayed in 96-well format, then screened for biofilm stimulation by ½ MIC cefixime in technical duplicate. Non-stimulated mutants were identified as having an average cefixime-induced biofilm of less than 200% of the vehicle control. Hits were validated in dose-response peg lid assays using cefixime, tobramycin, and thiostrepton and those lacking biofilm stimulation in response all three compounds were selected for follow up. The location of transposon insertions was identified by arbitrarily primed touchdown PCR followed by sequencing. B) A replicate plot showing biofilm stimulation by ½ MIC cefixime for the entire transposon library (∼13 000 mutants) across both technical replicates. The axes are shown as a log_10_ scale, and the dashed line shows the 200% of untreated control cutoff for hits. Each mutant is represented by a circle, with yellow circles showing the hits that failed to respond to three different antibiotics in dose-response follow-up assays. The identity of each verified hit is shown in the inset box. C) A schematic of the genes that, when disrupted, prevent biofilm formation in response to antibiotics is shown, with the genes of interest in different colours and neighbouring genes in grey. The predicted gene product description according to the *Pseudomonas* database^49^ for each hit is on the left.

### A conserved role for OprF in biofilm stimulation

Six mutants with diminished biofilm stimulation in response to all three antibiotics tested (**Figure S1**) were selected for further studies. *oprF* (*PA1777*) encodes the major outer membrane porin of *P. aeruginosa*. Two different mutants with insertions in *PA0163* were isolated; it encodes a predicted transcriptional regulator. Two mutants with independent insertions in *PA0177* were isolated; it encodes a CheW-like chemotaxis transducer from the Aer2 operon. *dsbA* (*PA5489*) encodes the thiol:disulfide interchange protein and *PA1895* encodes a predicted fatty acid desaturase. Two mutants with independent insertions in *PA2200* – encoding a predicted c-di-GMP phosphodiesterase – were also isolated (**Figure 3bc**).

We first focused on *oprF*, as it was implicated previously in *P. aeruginosa* biofilm formation.^50,51^ OprF is a highly abundant outer membrane porin, homologous to *E. coli* OmpA,^52^ and allows for the passive diffusion of small molecules across the outer membrane. OprF has important roles in *P. aeruginosa* virulence,^53^ immune sensing,^54^ and outer membrane structure.^55^ Loss of OprF is associated with membrane stress,^55^ as well as elevated levels of the secondary messenger c-di-GMP.^50^ We generated an *oprF* mutant in a clean background, confirmed that it lacked biofilm stimulation in response to sub-MIC antibiotics (**Figure 4a**), and complemented the stimulation phenotype with expression of the wild-type allele *in trans* (**Figure 4b**). Notably, the *oprF* mutant produced ∼2.5x more baseline biofilm than the wild type but levels did not further increase in response to antibiotic exposure, despite its wild type MICs for cefixime and tobramycin. We tested the *oprF* mutant with additional antibiotics, with different mechanisms of action: carbenicillin, chloramphenicol, ciprofloxacin, novobiocin, and trimethoprim. All stimulated biofilm formation of the PAO1 parent strain to varying degrees, with maximal stimulation at ¼ - ½ MIC; however, the *oprF* mutant failed to respond to any antibiotic with increased biofilm (**Figure 4c**). OprF has two conformations, closed and open. The closed conformation has a small N-terminal beta-barrel and a C-terminal peptidoglycan (PG)-binding domain, while the open conformation folds into a large beta-barrel. Interestingly, expression of a truncated version of OprF lacking its C-terminal PG-binding domain complemented the biofilm phenotype of the *oprF* mutant (**Figure 4d**), suggesting that this region is dispensable for biofilm stimulation. As *E. coli* is also capable of biofilm stimulation by sub-MIC antibiotics, we tested whether loss of *ompA*, the *oprF* homologue, similarly prevented biofilm stimulation. Cefixime, novobiocin, and tetracycline are potent biofilm-stimulating antibiotics for *E. coli* K12, and similar to *P. aeruginosa oprF*, the *ompA* mutant lacked antibiotic-induced biofilm stimulation (**Figure 4e**). This suggests a conserved role for OprF and OmpA in the response to sub-MIC antibiotics.

**Figure 4:**
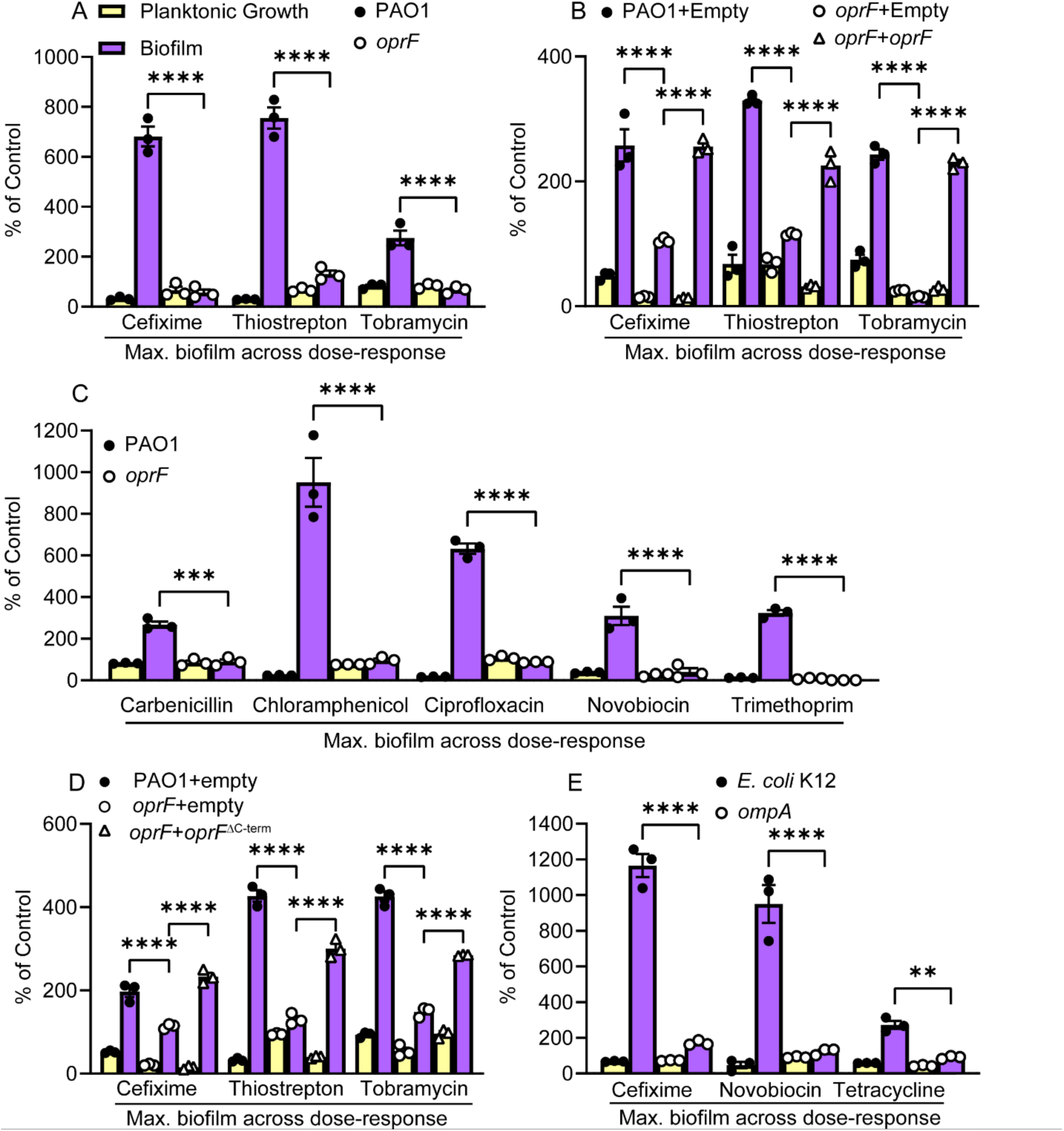
OprF is required for sub-MIC antibiotic-induced biofilm stimulation. A) Sub-MIC antibiotics fail to stimulate biofilm formation in an *oprF::FRT* mutant (N=3) B) Expression of OprF *in trans* restores the biofilm response to sub-MIC cefixime, thiostrepton, and tobramycin (N=3). C) Multiple antibiotics fail to stimulate biofilm formation of an *oprF* mutant (N=3). Notably, the *oprF* mutant was ∼16x more sensitive than the wild type to trimethoprim. D) Expression of a truncated OprF that lacks the C-terminal PG-binding region (*oprF*^ΔC-term^) restores antibiotic-induced biofilm formation in an *oprF* mutant (N=2). E) OmpA, the OprF homologue in *E. coli*, is required for biofilm stimulation by sub-MIC cefixime, novobiocin, or tetracycline in *E. coli* K12 (N=2). Planktonic growth (OD_600_, yellow) and biofilm (A_600_, purple) are reported as percentage of the vehicle control. A two-way ANOVA followed by Tukey’s multiple comparisons test was performed for the biofilm formation values for each panel. * = p<0.05, ** = p<0.01, *** = p<0.001, **** = p<0.0001. Each biological replicate has 3 technical replicates, and a representative biological replicate is shown with the circle or triangle symbols representing individual data points. The graphs show the maximum amount of biofilm observed from a dose-response assay (such as those shown in Figure 1) for each antibiotic/strain. Maximum biofilm formation occurred at 5 µM for cefixime, 10 µM for thiostrepton, 0.2 µM for tobramycin, 200 µM carbenicillin, 4 µg/mL for chloramphenicol, 0.25 µg/mL ciprofloxacin, 150 µg/mL novobiocin, and 16 µg/mL trimethoprim. For *oprF* mutant strains that are not complemented, maximum biofilm formation tended to occur at the lowest concentration of antibiotic as biofilm formation decreased at higher antibiotic concentrations. Error bars represent the standard error of the mean.

### A *sigX* mutant is deficient in the biofilm response to sub-MIC antibiotics

Prior studies linked regulation of OprF expression to the ECF sigma factor SigX in *P. aeruginosa*. When *oprF* is deleted, SigX activity increases.^50^ SigX controls the transcription of more than 250 genes, and *sigX* is immediately upstream of *oprF*.^56,57^ ECF sigma factors typically respond to environmental stimuli.^58^ The specific signals to which SigX responds are not well defined in *P. aeruginosa*, although it was linked to outer membrane stress and osmotic shock responses.^59^ Due to these predicted roles and its link to OprF expression, we hypothesized that *sigX* may be important for the biofilm response to sub-MIC antibiotics.

*sigX* was not a hit in our initial screen, so we generated a Δ*sigX* mutant and tested it for biofilm stimulation by sub-MIC cefixime, tobramycin, and thiostrepton. Like *oprF*, *sigX* failed to respond to these antibiotics (**Figure 5a**). The Δ*sigX* mutant displayed about 4.5x higher baseline levels of biofilm formation compared to the parent strain, but those levels did not increase further upon antibiotic treatment. The ECF sigma factor AlgU also regulates expression of *oprF*.^59^ We found that biofilm stimulation was unaffected in an *algU* mutant (**Figure S2**), so altering *oprF* regulation is insufficient to block biofilm stimulation by antibiotics. Fléchard *et al.* reported that Tween80 supplementation could restore wild-type membrane fluidity and gene expression in a *sigX* mutant.^60^ Addition of 0.1% Tween80 restored biofilm stimulation in a *sigX* mutant, but not in an *oprF* mutant (**Figure 5b**). Therefore, loss of *sigX* may disrupt biofilm stimulation indirectly due to defects in membrane fluidity.

**Figure 5:**
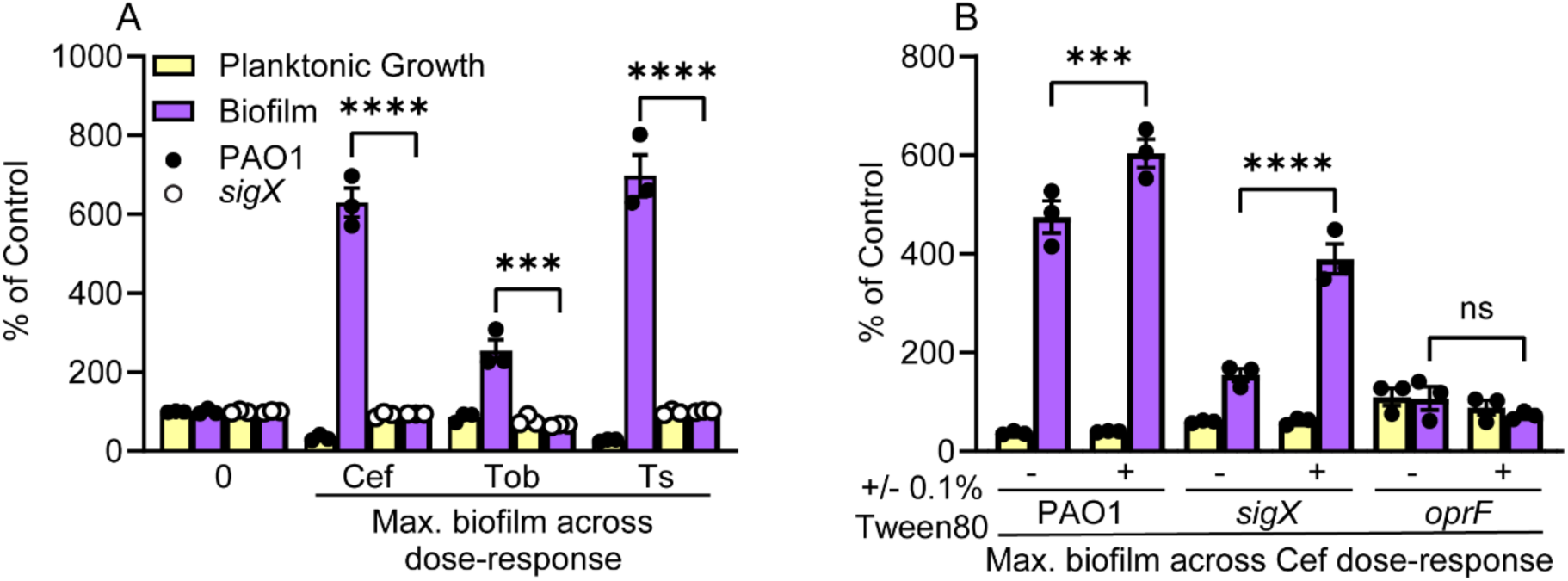
Sub-MIC antibiotics fail to stimulate biofilm formation in a *sigX* mutant. A) Sub-MIC cefixime, tobramycin, and thiostrepton do not stimulate biofilm formation in a *sigX* mutant (N=3). Notably, loss of *sigX* greatly increases the baseline biofilm formation in the untreated control. A two-way ANOVA followed by Šídák’s multiple comparisons test was performed between the biofilm formation data for PAO1 and the *sigX::FRT* mutant for each antibiotic in (A). Each data point is shown at individual circles, where black circles represent PAO1 and white circles represent *sigX::FRT*. Cef=cefixime, Tob=tobramycin, Ts=thiostrepton. B) Supplementing 0.1% Tween 80 into the media restores biofilm stimulation for a *sigX* mutant but not an *oprF* mutant (N=2). A two-way ANOVA followed by Tukey’s multiple comparisons test was performed for the biofilm formation data in (B). ns = not significant, * = p<0.05, ** = p<0.01, *** = p<0.001, **** = p<0.0001. For both figures, three technical replicates were performed for each biological replicate and data from a representative biological replicate is shown. The graphs show the maximum amount of biofilm observed from a dose-response assay (such as those shown in Figure 1) for each antibiotic/strain. Planktonic growth (OD_600_, yellow) and biofilm (A_600_, purple) are reported as percentage of the untreated control. Error bars represent the standard error of the mean.

### Probing potential links between disulfide bond formation and biofilm stimulation

OprF contains four cysteines that – depending on their redox state – affect its conformational bias (open versus closed).^61^ In considering how OprF may fit with our other hits, we noted that PA2200 has a periplasmic regulatory CSS domain that likely modulates phosphodiesterase activity in response to changes in disulfide bond formation, akin to the mechanism reported for the c-di-GMP phosphodiesterase PdeC from *E. coli*.^62^ CSS domains are characterized by a <u>c</u>ysteine-<u>s</u>erine-<u>s</u>erine motif that forms intramolecular disulfide bonds with another cysteine; in PdeC, formation of a disulfide bond by the CSS domain inhibits phosphodiesterase activity. Arr, a *P. aeruginosa* c-di-GMP phosphodiesterase that was previously implicated in biofilm stimulation by tobramycin, also has a periplasmic CSS domain.^34^ Further, disulfide bond formation requires DsbA, which was among the hits in our screen. To probe the role of disulfide bonds in biofilm stimulation, we performed biofilm assays using medium supplemented with dithiothreitol (DTT) at concentrations that did not impact growth. DTT directly reduces disulfide bonds to free thiols and suppresses disulfide bond formation. DTT supplementation suppressed biofilm stimulation by cefixime, thiostrepton, and tobramycin (**Figure 6a, b, and c**).

**Figure 6:**
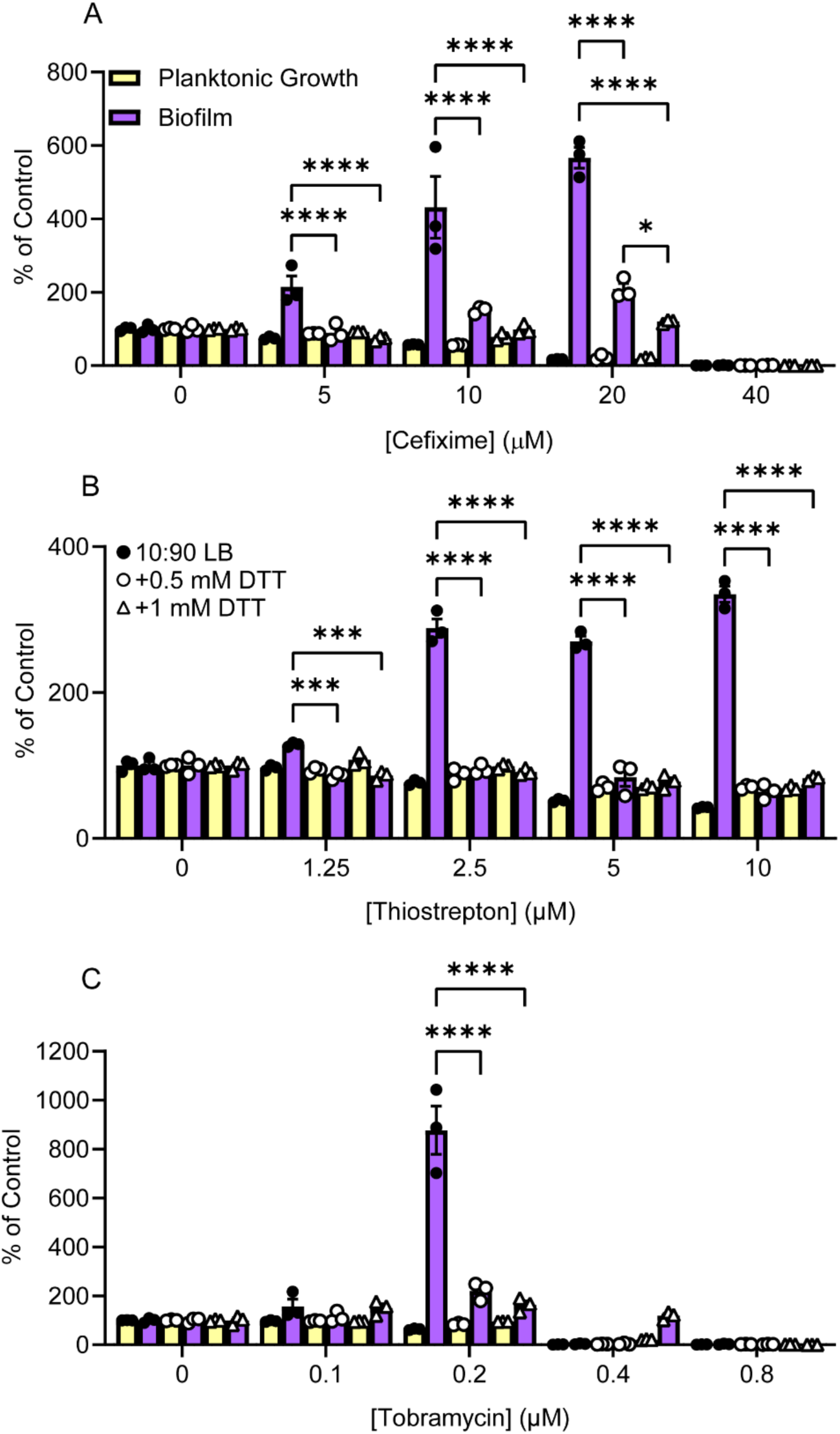
DTT suppresses biofilm stimulation by cefixime, thiostrepton, and tobramycin. Biofilm stimulation peg-lid assays were performed for A) cefixime, B), thiostrepton, or C) tobramycin in either 10:90 LB (black circles), or 10:90 LB with 0.5mM DTT (white circles) or 1mM DTT (white triangles) (N=2). A two-way ANOVA followed by Tukey’s multiple comparisons test was performed for the biofilm formation data in (B). * = p<0.05, ** = p<0.01, *** = p<0.001, **** = p<0.0001. Three technical replicates were performed for each biological replicate and data from a representative biological replicate is shown. Planktonic growth (OD_600_, yellow) and biofilm (A_600_, purple) are reported as percentage of the untreated control. Error bars represent the standard error of the mean.

*PA2200* is required for biofilm stimulation, and Arr, another c-di-GMP phosphodiesterase, was previously implicated in tobramycin-induced biofilm formation in a subset of *P. aeruginosa* strains, including PAO1.^34^ To determine if changes in c-di-GMP concentrations occur upon treatment with sub-MIC antibiotics, we used a reporter construct with the c-di-GMP responsive *cdrA* promoter driving expression of the *lux* operon.^63–65^ Treatment with sub-MIC thiostrepton and tobramycin significantly increased expression from the *cdrA* promoter compared to the vehicle control, whereas treatment with cefixime did not (**Figure 7a, b, and c**), although cefixime-treated cells showed a similar upward trend beginning at 10-12 hours. This suggests that c-di-GMP levels increased during antibiotic-induced biofilm formation in response to thiostrepton or tobramycin. In our prior experiments, introduction of a plasmid reduced biofilm stimulation by cefixime more so than thiostrepton or tobramycin, which could explain the lack of a signal increase in response to cefixime. In the *oprF* mutant, baseline *cdrA* promoter activity was higher than the wild-type – similar to previous reports – however, treatment with sub-MIC antibiotics did not further increase *cdrA* promoter activity over the vehicle control. (**Figure 7a, b, and c**).

**Figure 7:**
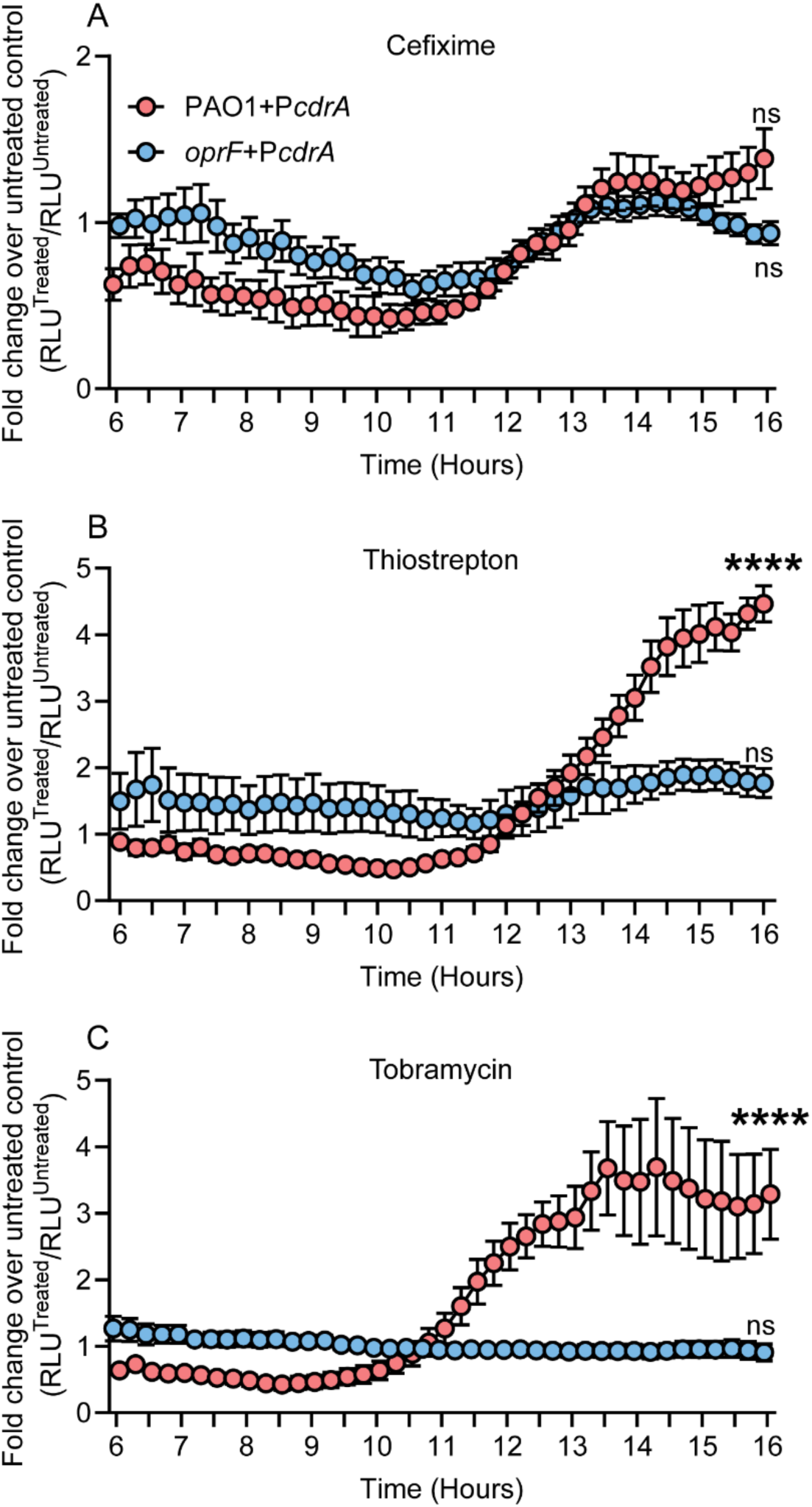
*cdrA* promoter activity increases following sub-MIC antibiotic treatment. PAO1 or *oprF::FRT* cells containing pMS402(Empty) or pMS402(P*cdrA*) were treated with ½ MIC cefixime (A), thiostrepton (B), tobramycin (C) or a vehicle control and monitored for growth and luminescence. Mean relative luminescence (RLU = arbitrary luminescence values divided by OD_600_) of the treated wells divided by the mean RLU of the matched untreated wells is plotted on the Y-axis as the fold change. The time in hours is plotted on the X-axis and readings were taken every 15 minutes. Circles represent the mean of six values taken across three biological replicates each performed in technical duplicate. Error bars represent the standard error of the mean. Red and blue circles represent data from PAO1 and *oprF::FRT* containing pMS402(P*cdrA*), respectively. Data are plotted as a fold change over the empty vector control, therefore the empty vector data are all equal to one, as they would be a well’s fold change versus itself. A two-way ANOVA followed by Dunnett’s multiple comparisons test was performed to compare each antibiotic treatment condition across the time course to the six-hour time point, and the results of the comparison are shown for the 16-hour time point. ns = not significant, **** = p<0.0001.

## Discussion

A variety of antibiotics with different modes of action can stimulate *P. aeruginosa* biofilm formation at subinhibitory concentrations. We first tested the idea that they kill a susceptible subpopulation of cells, releasing resources such as eDNA that can then seed biofilm development by the remaining cells. This effect was reported for *Enterococcus faecalis* and *Haemophilus influenzae* treated with antibiotics targeting peptidoglycan synthesis. Other classes of antibiotics, such as protein synthesis inhibitors or fluoroquinolones, did not induce biofilm formation.^38,39^ In contrast, sub-MIC fluoroquinolones induce self-lysis via activation of prophage elements in *P. aeruginosa*, releasing eDNA.^41^ We found that adding genomic eDNA, cell lysate, or a rapidly lytic antibiotic (polymyxin B) failed to phenocopy the amount of biofilm observed after exposure to sub-MIC antibiotics, suggesting the response is more complex. The lack of biofilm stimulation by eDNA could be due to its distribution in the medium instead of concentration at the substratum.^66^ A limitation of using purified genomic DNA is the loss of DNA-associated proteins that could be important for adhesion.^67^ To circumvent that possibility, we also tested whole-cell lysate, but only saw minor increases in biofilm at higher lysate concentrations. If cell lysis drove the biofilm response to antibiotics, we might expect less response to bacteriostatic versus bactericidal antibiotics. Instead, we saw high levels of biofilm stimulation with sub-MIC chloramphenicol and thiostrepton, and comparatively weak biofilm stimulation with polymyxin B.

These data led us to hypothesize instead that there is a coordinated response to antibiotic exposure, and we used a transposon mutant screen to identify genes involved in those pathways. Analysis of our screening hits suggested that biofilm stimulation may be driven in part by antibiotic stress-induced changes in cysteine redox state in the periplasm. For example, distribution of OprF between its open and closed conformation is modulated by disulfide bonds catalyzed by DsbA. The open conformation is more abundant when cysteine to serine mutations are introduced in the residues that form two disulfide bonds.^61^ Loss of *dsbA* inhibited biofilm stimulation by sub-MIC antibiotics. Similar to some of our other hits, the *dsbA* mutant had elevated basal biofilm formation compared to the parent strain, but failed to increase further upon antibiotic exposure. This phenotype suggests that loss of these gene products could pre-activate the stress pathways that lead to increased biofilm formation. Prior work in *P. aeruginosa* showing that loss of *dsbA* increased c-di-GMP levels in a WspR-dependent manner following treatment with cell wall-active antibiotics.^68^ *P. aeruginosa dsbA* is in a predicted operon with the diguanylate cyclase gene *dgcH*, indicating potential co-regulation of c-di-GMP levels with disulfide bond formation.

Our previous work on antibiotic-induced biofilm formation in *E. coli* showed that the response involved changes in NADH redox state and was abrogated by disrupting central metabolism and respiration pathways.^69^ Periplasmic disulfide bond formation liberates electrons from free thiols that are shuttled through DsbA and DsbB to respiratory quinones, which then participate in the electron transport chain.^70^ Our *E. coli* biofilm stimulation work also implicated ArcB and NlpE, which are involved in signal transduction and whose activity is regulated by disulfide bonding.^71,72^ A *dsbA* mutant of *E. coli* failed to respond to sub-MIC novobiocin, and narrowly missed our arbitrary hit cutoff for cefixime and tetracycline.^69^ Since antibiotics alter respiratory chain flux,^73^ and altering respiration activity modulates antibiotic-induced biofilm formation,^69^ it is reasonable to speculate that the ability of DsbA/B to cycle electrons to respiratory quinones could also be disrupted by sub-MIC antibiotics.

The importance of normal disulfide bonding was confirmed by the lack of biofilm stimulation in the presence of DTT, which causes widespread disruption of disulfide formation. Undoubtably, cysteines in proteins other than our hits are affected, however growth was not affected at the concentrations we used. To more directly probe the involvement of the Dsb system, future work could use a recently discovered DsbB inhibitor that is more specific than DTT. ^74,75^ The emerging model from our studies in both *P. aeruginosa* and *E. coli* suggests that antibiotic-induced biofilm formation occurs at a crossroads between energetics and periplasmic redox state.

Of the regulators of *oprF* expression, we showed that SigX but not AlgU is important for biofilm stimulation. The molecular cues for SigX activation are unknown, although its activity increases in an *oprF* background.^50^ We speculate that antibiotic stress affects folding of OprF and integration of the N-terminal domain into the outer membrane, which in turn disrupts integrity of the envelope, leading to activation of SigX. If SigX coordinates a cell envelope stress response,^58^ then OprF misfolding could cause significant envelope stress, as OprF is a highly abundant outer membrane protein and important for envelope integrity.^76^ Alternatively, OprF could be involved in surface sensing and act as a checkpoint for initiating biofilm formation when under antibiotic stress. In *Salmonella*, redox stress drives conformational changes in OmpA by impacting disulfide bond formation.^77^ In *E. coli*, OmpA interacts with NlpE and is required for NlpE-mediated induction of the Cpx periplasmic stress response upon surface adhesion.^78^ Previous work in *E. coli* also implicated OmpA in mediating biofilm formation via Cpx.^79^ Despite having the core components of the Cpx system, *P. aeruginosa* lacks an obvious *nlpE* homologue. It is unclear if the *P. aeruginosa* Cpx system is involved in biofilm stimulation, or if SigX fills this role.

We also identified the putative c-di-GMP phosphodiesterase PA2200 as important for biofilm stimulation. PA2200 has a periplasmic, N-terminal CSS domain that likely regulates its PDE activity in response to cysteine redox state, similar to *E. coli* PdeC.^62^ Hoffman *et al.* identified another PDE, Arr, as an important player in the biofilm response to tobramycin.^34^ *arr* is not present in all *P. aeruginosa* strains capable of the biofilm response,^35^ and is less conserved in *P. aeruginosa* than *PA2200*. The presence of *PA2200* could explain how strains lacking *arr* can still undergo biofilm stimulation. According to the *Pseudomonas* database, there are 350 *PA2200* orthologs, but only 64 *arr* orthologs.^49^ The redundancy of *arr* and *PA2200* in the PAO1 genome suggests potential specialization of these two genes. They could be differentially regulated, or control the concentrations of spatially distinct, local c-di-GMP pools in the cell. Indeed, in response to sub-MIC tobramycin treatment, the promoter activity of *arr* and *PA2200* increase and decrease, respectively.^80^ Using a reporter construct where c-di-GMP-bound FleQ upregulates expression from the *cdrA* promoter,^63–65^ we showed that c-di-GMP levels increased upon treatment with sub-MIC thiostrepton and tobramycin. There is an apparent contradiction in the requirement for a c-di-GMP phosphodiesterase in a response that entails increases in c-di-GMP levels. Hoffman *et al.* ascribe this discrepancy to the wide range of effects that inactivating individual DGCs or PDEs has on biofilm formation.^34^ We suggest that the biofilm stimulation response may require temporal regulation of c-di-GMP, where lower c-di-GMP is required in the early stages of the response pathway. Indeed, we saw a signal increase in *cdrA* promoter activity only after 10-12 hours of growth of PAO1 in sub-MIC thiostrepton or tobramycin.

An advantage of the *cdrA* promoter reporter over exogenous, synthetic reporter systems is that *cdrA* is native to *P. aeruginosa*. While we used this reporter as a proxy for c-di-GMP levels, we can also conclude that sub-MIC thiostrepton and tobramycin induce expression of CdrA, an adhesin that is important for cell-cell and cell-matrix interactions in biofilms.^81^ While more work is needed to confirm that levels of functional, surface-exposed CdrA increase, our work provides preliminary evidence that increases in CdrA may be a factor in antibiotic-induced biofilm formation.

In addition to *oprF*, *PA2200*, and *dsbA*, we identified *PA0177*, *PA0163*, and *PA1895* as important for biofilm stimulation. *PA0177* is part of the *aer2* aerotaxis operon that signals in the presence of oxygen,^82^ and is upregulated upon surface attachment.^83^ *PA0163* is a predicted but poorly characterized AraC-type transcriptional regulator. *PA1895* is part of the QscR-regulated operon that delays induction of the LasR and RhlR quorum sensing regulon.^84^ Apart from *PA0163*, the remaining uncharacterized hits may impact biofilm formation through quorum sensing (*PA1895*) or via regulatory changes in response to electron acceptor availability (*PA0177*). Notably, our screen identified no mutants deficient in EPS biosynthesis or surface attachment, such as *pelD* or *sadC*. This could be due to redundancy in matrix components, or to the ability of even poor biofilm formers to be sufficiently responsive to sub-MIC antibiotic exposure that they passed our 200% increase cut-off.

Overall, this work implicates changes to disulfide bonding and envelope stress signalling in the biofilm stimulation response. Based on our studies in *E. coli* and *P. aeruginosa*, we propose a testable model where: 1) antibiotic stress affects flux through central metabolism and the electron transport chain due to the energy demand of damage repair; 2) perturbing flux through the electron transport chain affects the ability of the periplasmic disulfide bonding system to replenish DsbA by shunting electrons to respiratory quinones; 3) disrupting periplasmic cysteine redox homeostasis activates signalling through sentinel proteins such as NlpE, ArcB, and OmpA in *E. coli*, or PA2200/Arr and OprF in *P. aeruginosa*; and 4) these sensors and sentinels activate signalling pathways, such as those impacting c-di-GMP metabolism, that induce biofilm formation. While our data are in line with this model, it must be further tested.

Biofilm stimulation occurs in response to both bacteriostatic and bactericidal antibiotics, which suggests that either independent types of stressors converge on the same phenotype through different pathways, or that distinct stressors have a common impact on the cell that triggers shared pathways. Our prior studies in *E. coli* suggested that bactericidal and bacteriostatic antibiotics drive biofilm formation through opposing effects on energy metabolism.^69^ The biofilm stimulation response could also have evolved under pressure from different kinds of stress, such as phage infection or interbacterial contact-dependent antagonism,^66^ and antibiotic stress merely taps into this established phenotype. We also considered the idea that if antibiotics are signalling molecules,^85^ they could act as cues for the establishment of a cooperative multispecies biofilm.^86^ Antibiotics in the environment could signal the boundaries of a “territory” of producers, where *P. aeruginosa* reacts by forming a biofilm at a sub-MIC “border”. Whatever the true role of antibiotics is in nature, it is clear that the bacterial response to antibiotics is integrated deeply into their biology.

## Materials and Methods

### Bacterial Strains and culture conditions

All strains (listed in table S1) were stored in 15% glycerol stocks at -80°C and frozen stocks were used to inoculate overnight cultures, which were grown at 37°C, 200 rpm. Ninety-six well plates were incubated in humidified containers to prevent evaporation in peripheral wells. Lysogeny Broth (LB) media (Bioshop) contained 10g/L tryptone, 5g/L yeast extract, and 5g/L sodium chloride. The 10% LB-PBS media (10:90 LB) was made by diluting LB 1:9 with 1x phosphate buffered saline (PBS). Apart from polymyxin B and ciprofloxacin, which were solubilized in sterile milliQ water, all compounds used for biofilm assays were solubilized in DMSO or water and diluted in growth media such that the final concentration of DMSO never exceeded 1.33% (v/v). Antibiotic selection was performed where appropriate with the following concentrations: 15 or 30 µg/mL gentamicin (Gent15/30) for *E. coli* and *P. aeruginosa*, respectively; 50 or 150 µg/mL kanamycin (Kan50/150) for *E. coli* and *P. aeruginosa*, respectively; 100 µg/mL ampicillin (Amp100) or 200 µg/mL carbenicillin for *E. coli* and *P. aeruginosa*, respectively. In experiments where pHERD30T was used for gene expression, sterile L-arabinose was added at a final concentration of 0.1% (v/v) to induce promoter activity.

### Antibiotic-induced biofilm formation assays

Antibiotic-induced biofilm formation assays were performed as previously described with modifications.^87^ PAO1 strains were cultivated overnight at 37°C, 200 rpm in 10:90 LB media. Overnight cultures were diluted 1:25 in 10:90 LB media and sub-cultured to OD_600_ = 0.1 under the same growth conditions. Subcultures were then diluted 1:500 in fresh 10:90 LB. Assays were prepared in 96-well plates with 96-peg lids (Nunc). Wells contained 150 µL of total culture, with 148 µL of the diluted subculture added to each well (sterility control wells contained 148 µL of media in place of the subculture) and 2 µL of either an antibiotic or DMSO. Antibiotic-treated wells contained 2 µL of antibiotic suspended in DMSO or sterile water, while the vehicle and sterility control wells contained 2 µL DMSO or water. Assays were incubated at 37°C, 200 rpm for 16 hours in humidified containers. For PA14 and *E. coli* strains, the procedure is identical except that cells were grown in 50:50 LB (one to one ratio of LB and 1x PBS). Peg lids were removed from the plates and planktonic growth was measured by reading optical density at 600 nm. Peg lids were submerged in 1x PBS for 10 min to remove loosely attached cells, then transferred to 0.1% crystal violet for 15 min to stain adhered cells. Peg lids were removed from crystal violet and washed immediately by submerging in 70 mL milliQ water in a basin, then transferred to a fresh milliQ water basin for 10 min. Three additional 10-min washes with milliQ water were performed in succession to remove excess stain. After washing, peg lids air dried for a minimum of 30 min. Stained biofilms were solubilized in 200 µL of 33.3% acetic acid in a 96-well plate for 5 min. The absorbance of the eluted crystal violet dye was quantified at 600 nm using a plate reader. Optical density (planktonic growth) and absorbance values (biofilm) were plotted as the percent of the vehicle control values (corrected for the sterility well background).

### Creation of a PAO1 Himar1 Mariner transposon library

Transposon mutagenesis was performed as previously described, with modifications.^88^ *E. coli* SM10 λ pir cells were transformed with pBT20 (carrying the Himar1 Mariner transposon) to create *E. coli* SM10 λ pir /pBT20. Successful transformants were selected with Amp100 on solid media. *P. aeruginosa* PAO1 was grown on an LB agar plate overnight and *E. coli* SM10 λ pir /pBT20 was grown on an Amp100 LB agar plate overnight at 37°C. A full inoculating loop of cells was scraped from each plate and resuspended in either 1 mL of LB media (for PAO1) or 1 mL of LB Amp100 (for *E. coli* SM10 λ pir /pBT20). Five hundred microlitres of the *E. coli* SM10 λ pir /pBT20 and 500 µL of PAO1 were mixed together in a new tube and centrifuged for 1 min at 21 000 x g to pellet the cells. Eight hundred microlitres of supernatant was removed and the mixed cell pellet was resuspended in the remaining supernatant. A mating spot was created by placing 100 µL of the resuspended mixed cell pellet in a single spot in the middle of an LB agar plate. The mating spot was dried at room temperature for 20 min and the plate incubated at 37°C overnight. The mating spot was collected using a sterile loop and resuspended in 1 mL of LB. *P. aeruginosa* PAO1 transposon mutants were selected by plating 100 µL of the mating spot cell suspension on *Pseudomonas* Isolation Agar (PIA) (BD Difco) + 100 µg/mL gentamicin. PIA contains 25 µg/mL irgasan, which selects against the *E. coli* donor. Single colonies were picked and arrayed into 96-well plates containing 100 µL of Gent30 LB. On each plate, six wells containing LB media per 96-well plate were inoculated with the parental strain PAO1 (wells H1-6) and another six wells containing LB media were left blank as sterility controls (H7-12). The 96-well plates were incubated overnight at 37°C and 200 rpm in humidified containers. After incubation, 100 µL of LB + 30% glycerol was added to each well of the 96-well plates and the plates were stored at -80°C. We collected a total of 165 96-well plates, totaling 13 776 mutants.

### Screening PAO1 transposon mutants for antibiotic-induced biofilm formation

The screening protocol was developed based on the antibiotic-induced biofilm formation assay. *P. aeruginosa* PAO1 transposon mutants were inoculated from 15% glycerol freezer stocks into a 96-well plate containing 10:90 LB media (150 µL/well) using a 96-pin tool. The plates were incubated overnight at 37°C, 200 rpm in a humidified container. A 96-well subculture plate containing 10:90 LB media (144 µL/well) was then inoculated with 6 µL of overnight culture. The subculture plate was incubated at 37°C while shaking at 200 rpm for 2 hours. Using the subculture plate and a 96-pin tool, we inoculated three assay (96 peg lid) plates containing 148 µL of 10:90 LB with two plates containing 2 µL of cefixime (1/2 MIC final concentration) and one plate containing 2 µL DMSO (1.33% v/v final). Sterility and vehicle control wells in row H contained 1.33% (v/v) DMSO instead of antibiotic. A polystyrene 96-peg lid (Nunc) was used as a substratum for biofilm growth. The plates were incubated overnight at 37°C, 200 rpm in humidified containers. The planktonic growth and biofilm for all plates were quantified as per the protocol in “Antibiotic-induced biofilm formation assays” (above). An *a priori* cut-off for biofilm production in wild type was defined as >200% increase in biofilm in the presence of sub-MIC cefixime when compared with the vehicle control, dimethyl sulfoxide (DMSO). Thus, hits from the screen were defined as mutants that failed to meet this cut-off (i.e. produced less than 200% of control biofilm in cefixime). Mutants that were hits in both replicates of the screening assay were selected for verification in a dose-response biofilm stimulation assay with cefixime, thiostrepton, and tobramycin.

### Transposon insertion site identification

The sites of transposon insertion in validated mutants were identified using a semi-random, two-step (ST)-PCR method.^89^ The first round of PCR was conducted with a Himar1 Mariner-specific primer (Himar1 Primer PCR Round 1) and a hybrid consensus-degenerate primer (Arbitrary Primer PCR Round 1). Random annealing of the consensus-degenerate primer was facilitated by starting with an annealing temperature of 65°C and reducing the temperature by 0.5°C for 30 cycles.^31^ The second round of PCR was conducted with a transposon primer (Himar1 Primer PCR Round 2) and a primer specific for the arbitrary primer used in the first round (Arbitrary Primer PCR Round 2) at an annealing temperature of 58°C for 40 cycles to ensure sufficient amplification. The second-round PCR products were gel-excised and purified using a GeneJET gel extraction kit (Thermo) followed by Sanger sequencing of the PCR fragments using the sequencing primer (TD PCR Sanger Sequencing Primer). Sequences were mapped to the *P. aeruginosa* PAO1 reference genome found at www.pseudomonas.com using the BLAST search function.^49^

### eDNA and cell lysate biofilm assays

Chromosomal DNA was isolated from *P. aeruginosa* PAO1 cells using a Promega Wizard Genomic DNA Purification Kit. Purified genomic DNA (gDNA) was resuspended to 10 ng/µL in nuclease-free water. Cell lysates were prepared using freeze-thaw cycling on *P. aeruginosa* PAO1 cells grown to OD_600_ = 1.8 in LB media. Seven hundred microlitres of 1.8 OD_600_ culture was incubated at - 80°C for 30 min and thawed at room temperature for 30 min. This freeze-thaw cycle was repeated at least five times for a total of at least six cycles. The freeze-thaw method resulted in >99% cell lysis, which was verified by CFU counts of cells pre- and post-lysis. Biofilm formation assays were set up as described in “Antibiotic-induced biofilm formation assays”, except 15 µL of gDNA or cell lysate (at the indicated concentrations) were added to each treated well, with 135 µL of bacterial subculture. For vehicle and sterility control wells, water was used in place of eDNA and LB was used in place of cell lysate. The assays were performed in 10:90 LB media at 37°C, 200 rpm for 16 hours. Biofilms were stained and analyzed as described above in “Antibiotic-induced biofilm formation assays”.

### Construction of *sigX*::FRT and *oprF*::FRT mutants

FRT mutants were constructed as previously described.^90^ Briefly, DNA spanning from *sigX* to the end of *oprF* was PCR amplified from PAO1 chromosomal DNA using the *sigX* Fwd and *oprF* Rvs primers. The amplified DNA was ligated into pEX18Ap at the EcoRI and HindIII cut sites. The resulting construct was then digested with EcoRV or SmaI, corresponding to native internal sites in *sigX* or *oprF*, respectively. A gentamicin cassette flanked by FRT sites (FRTGm) was excised from the pPS856 plasmid with EcoRV or SmaI and ligated into the corresponding cut site of pEX18Ap containing EcoRV or SmaI *sigX*-*oprF*. The assembled pEX18Ap(*sigX*-*oprF::*FRTGm) was moved into *E. coli* SM10, then mated into PAO1. PAO1 cells containing the single crossover were selected for on *Pseudomonas* Isolation Agar (PIA) containing 200 µg/mL carbenicillin. Colonies were streaked on LB (no NaCl) plates containing 5% sucrose and 30 µg/mL gentamicin to select for plasmid loss and a second recombination event. Single colonies were then patched onto plates containing 200 µg.mL carbenicillin or 30 µg.mL gentamicin, and colonies that only grew on gentamicin were selected for colony PCR to confirm the presence of the gentamicin cassette. To excise the gentamicin resistance cassette, pFLP2 was introduced into a colony containing the correct FRTGm insertion using electroporation. Cells containing pFLP2 were selected for on 200 µg/mL carbenicillin plates. To select for the loss of pFLP2, colonies from the carbenicillin plate were streaked onto sucrose plates. Colonies from the sucrose plates were patched onto sucrose and gentamicin/carbenicillin plates; cells that only grew on sucrose were presumed to have lost pFLP2 and the gentamicin cassette, which was validated by PCR.

### *cdrA* promoter reporter

The *cdrA* promoter region was cloned upstream of the *lux* genes on the plasmid pMS402 as described previously.^65^ Expression of *cdrA* activated by c-di-GMP bound FleQ is used as a proxy for cellular c-di-GMP levels.^63^ Overnight cultures of PAO1 or *oprF* cells containing pMS402(Empty) or pMS402(P*cdrA*) were made by inoculating 3 mL of LB+Kan150 from frozen stocks and were incubated at 37 °C with shaking at 200 rpm. Subcultures were made by transferring 120 µL of the overnights into 3 mL of 10:90 LB+Kan150 and incubated at 37 °C with shaking at 200 rpm for ∼3 hours until an OD_600_ of ∼0.1 was reached. Cultures were normalized to an OD_600_ of 0.1, then diluted 1:500 in fresh 10:90 LB+Kan150. Assays were prepared in white-walled 96-well plates with clear bottoms (Corning). Serial dilutions of cefixime, tobramycin, or thiostrepton were added to the 96-well plate in the same manner as the antibiotic induced biofilm formation assays. Afterwards, 148 µL of the diluted culture was added to each well of the plate, except row H, which served as a sterility control and only received 148 µL of 10:90 LB. The plate was incubated at 37 °C with continuous double orbital shaking for 16 hours in a Synergy Neo (Biotek) plate reader. Growth (OD_600_) and luminescence (luminescence fiber) measurements were taken every 15 minutes. Relative luminescence units (RLU) were calculated by dividing each well’s luminescence value by its growth at the corresponding time point. Relative changes in RLU compared to the untreated control were calculated by dividing each well’s RLU value by the untreated control matching the same plate column and timepoint.

### Data analysis and graphs

Data from plate readers were analyzed using Microsoft Excel and Graphpad Prism 9. All graphs were created using Prism 9. Statistical significance was determined by the procedure outlined in each figure caption. Sequencing data was analyzed using Geneious 6.0.6.

## Author contributions

LNY, MRMR, and LLB designed the study. LNY, MRMR, MR, SKP and JC generated the transposon library and screened it for non-responsive mutants. LNY, AMG, JC, and HH performed follow-up studies of selected hits. LNY, MRMR, and LLB wrote and edited the manuscript. All authors read and approved the final manuscript.

## Acknowledgements

We thank Michael Surette for the use of an automated colony picker and guidance in transposon library construction, and Uyen Nguyen and Neha Sharma for assistance in early development of the biofilm stimulation assay. This work was funded by the Natural Sciences and Engineering Research Council of Canada (www.nserc-crsng.gc.ca) grant RGPIN-2021-04237 to LLB. LLB holds a Tier I Canada Research Chair in Microbe-Surface Interactions. LNY holds an NSERC PGS-D award. MRMR held an Ontario Graduate Fellowship. SKP received an undergraduate research award from Glyconet. MR held an NSERC USRA scholarship.

## Data Availablility

All data generated and analysed during this study are included in this article and its supplementary information files.

