## Supplementary Table S1 and Supplementary Figures S1 and S2 for "A genetic screen identifies a role for *oprF* in *Pseudomonas aeruginosa* biofilm stimulation by subinhibitory antibiotics"

**Table S1: Strains and primers used in this study.**

| **Strain Name** | **Genotype/Characteristics** | **Reference** |
| --- | --- | --- |
| *Pseudomonas aeruginosa* |  |  |
|  | PAO1 wild type strain. Graciously donated by Keith |  |
| PAO1 | Poole (Queen's University, Kingston, Canada) | ^1^ |
| PAO1 *oprF::Himar1* | Himar1 transposon insertion mutant in *oprF*. Gentamicin resistant. Library ID: BBtn1_G3 | This Study |
| PAO1 *PA2200::Himar1* | Himar1 transposon insertion mutant in *PA2200*. Gentamicin resistant. Library ID: BBtn66_B3 and BBtn72_E12 | This Study |
| PAO1 *dsbA::Himar1* | Himar1 transposon insertion mutant in *dsbA*. Gentamicin resistant. Library ID: BBtn52_E12 | This Study |
| PAO1 *PA0177::Himar1* | Himar1 transposon insertion mutant in *PA0177*. Gentamicin resistant. Library ID: BBtn79_C6 and BBtn82_C3 | This Study |
| PAO1 *PA0163::Himar1* | Himar1 transposon insertion mutant in *PA0163*. Gentamicin resistant. Library ID: BBtn34_A9 BBtn82_F6 | This Study |
| PAO1 *PA1895::Himar1* | Himar1 transposon insertion mutant in *PA1895*. Gentamicin resistant. Library ID: BBtn74_A3 | This Study |
| PAO1 *oprF*::FRT | *oprF* FRT mutant with the Gentamicin cassette flipped out. | This Study |
| PAO1 *sigX*::FRT | *sigX* FRT mutant with the Gentamicin cassette flipped out. | This Study |
| PAO1 + pHERD30T | PAO1 with empty pHERD30T. Gentamicin resistant. | This Study |
| PAO1 *oprF*::FRT + pHERD30T | *oprF* FRT mutant with empty pHERD30T. Gentamicin resistant. | This Study |
| PAO1 *oprF*::FRT + pHERD30T-*oprF* | *oprF* FRT mutant expressing WT *oprF* from pHERD30T. Gentamicin resistant. | This Study |
| PAO1 *oprF*::FRT + pHERD30T-*oprF^trunc^* | *oprF* FRT mutant expressing *oprF* with a C-terminal truncation (residues 1-184 in the full-length polypeptide numbering) from pHERD30T. Gentamicin resistant. | This Study |
| PAO1 *oprF*::FRT + pHERD30T-*oprF* A312C | *oprF* FRT mutant expressing *oprF* with an A312C mutation (numbering referrers to mature polypeptide. A312 is A336 in full-length polypeptide). Gentamicin resistant. | This Study |
| PAO1 + pMS402 | PAO1 with promoter-less pMS402, which contains the *luxCDABE* operon. Kanamycin resistant. | ^2^ |
| PAO1 + pMS402-P*cdrA* | PAO1 with pMS402 containing the *cdrA* promoter in front of the *luxCDABE* operon. Kanamycin resistant. | ^2^ |
| PAO1 *oprF*::FRT + pMS402 | PAO1 *oprF* FRT mutant with promoter-less pMS402, which contains the *luxCDABE* operon. Kanamycin resistant. | This Study |
| PAO1 *oprF*::FRT + pMS402-P*cdrA* | PAO1 *oprF* FRT mutant with pMS402 containing the *cdrA* promoter in front of the *luxCDABE* operon. Kanamycin resistant. | This Study |
| PA14 | PA14 wild type strain. | ^3^ |
| PA14 *algU* | PA14 containing an *algU* clean deletion. | ^2^ |
| *Escherichia coli* |  |  |
| DH5α | F– *endA1 gln*V44 *thi*-1 *recA1 relA1 gyrA96 deoR nupG purB20 φ80dlacZΔM15 Δ(lacZYA-argF)U169*, *hsdR17(rK–mK+)*, λ–. Used for amplifying plasmid DNA and transformations. | Invitrogen |
| DH5α + pHERD30T | DH5α containing pHERD30T. Gentamicin resistant. | This Study |
| DH5α + pHERD30T-*oprF* | DH5α containing pHERD30T-*oprF*. Gentamicin resistant. | This Study |
| DH5α + pMS402 | DH5α containing pMS402. Kanamycin resistant. | ^2^ |
| DH5α + pMS402-P*cdrA* | DH5α containing pMS402-P*cdrA*. Kanamycin resistant. | ^2^ |
|  | Used for efficient conjugative transfer of plasmid DNA to |  |
| SM10-λ*pir* | *P. aeruginosa*. | ^4^ |
|  | SM10 containing *λpir* which allows for replication of |  |
|  | plasmids with *oriR6K* origins. Used to transfer pBT20 to |  |
| SM10-λ*pir* + pBT20 | *P. aeruginosa* PAO1. | This Study |
| **Primer Name** | **Sequence (5’ to 3’)** | |
| +*oprF* Fwd | GTACGAATTCGATGGGGATTTAACGGATG | |
| +*oprF* Rvs | GCATAAGCTTGCTCAGCCGATTACTTG | |
| +*sigX* Fwd | CTGAGAATTCGCACTCGGAGCTGTTCCAC | |
| +*oprF^trunc^* Rvs | TATAAAGCTTTTAGAAGTTGAAGCCGA | |
| +*ompA* Fwd | TATAGAATTCTGGCGTATTTTGGATGATAACGAGGC | |
| +*ompA* Rvs | ATAGAAGCTTGTTTTTCTACCAGACGAGAACTTAAGC | |
| Arbitrary Primer PCR Round 1 | GGCCACGCGTCGACTAGTACNNNNNNNNNNAGAG | |
| Himar1 Primer PCR Round 1 | TATAATGTGTGGAATTGTGAGCGG | |
| Arbitrary Primer PCR Round 2 | GGCCACGCGTCGACTAGTAC | |
| Himar1 Primer PCR Round 2 | ACAGGAAACAGGACTCTAGAGG | |
| TD PCR Sanger Sequencing Primer | CACCCAGCTTTCTTGTACAC | |

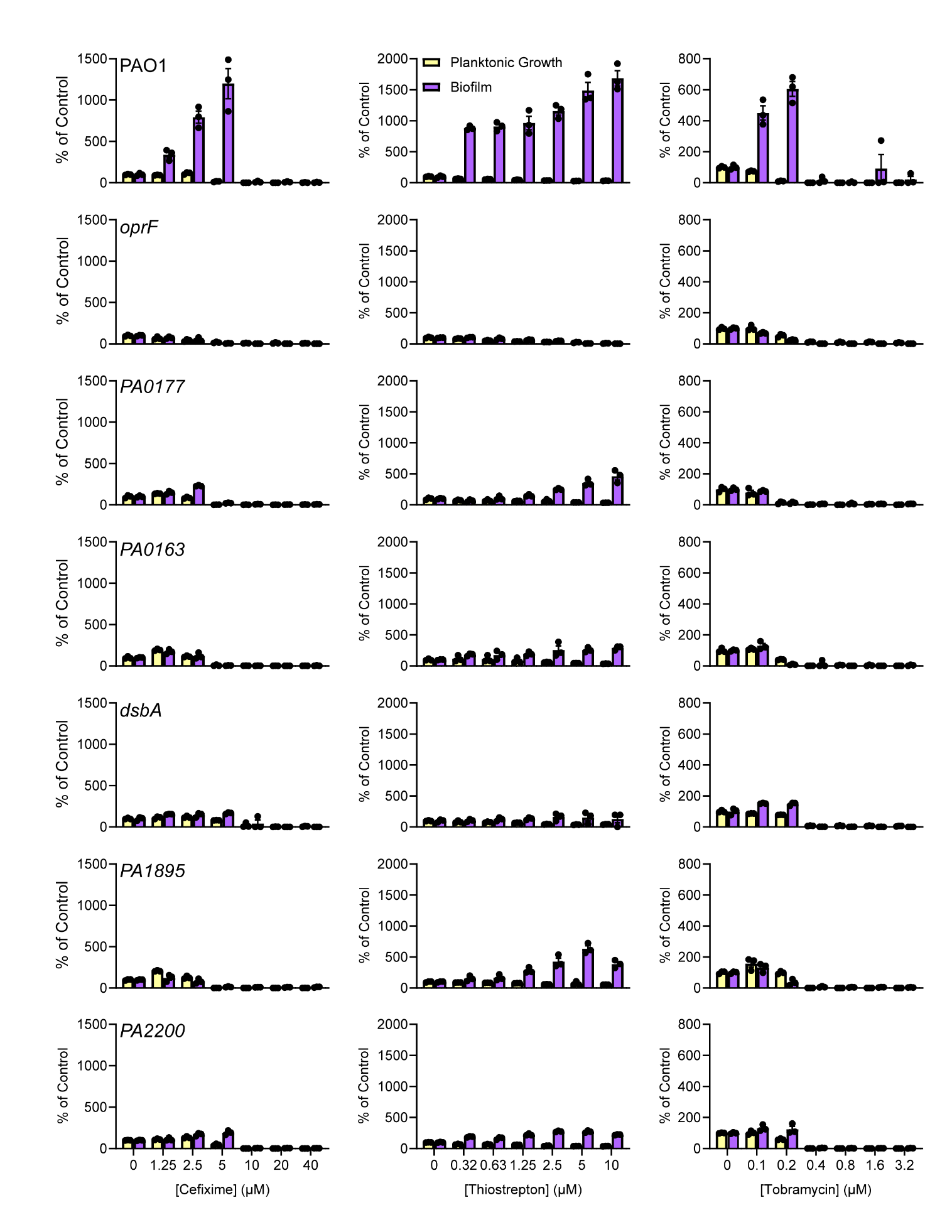

**Figure S1: Mutants unable to respond to three different sub-MIC antibiotics.** Cefixime (left column), thiostrepton (middle column), and tobramycin (right column) induce biofilm formation in PAO1 (top row), but not the transposon mutant screen hits (remaining rows). The gene name of each mutant is indicated above the cefixime graph for each respective row. Planktonic growth (OD_600_, yellow) and biofilm (A_600_, purple) are reported as percentage of the untreated treated control. Two biological replicates were performed with 3 technical replicates for each, and a representative biological replicate is shown with the circles representing individual data points. Error bars represent the standard error of the mean.

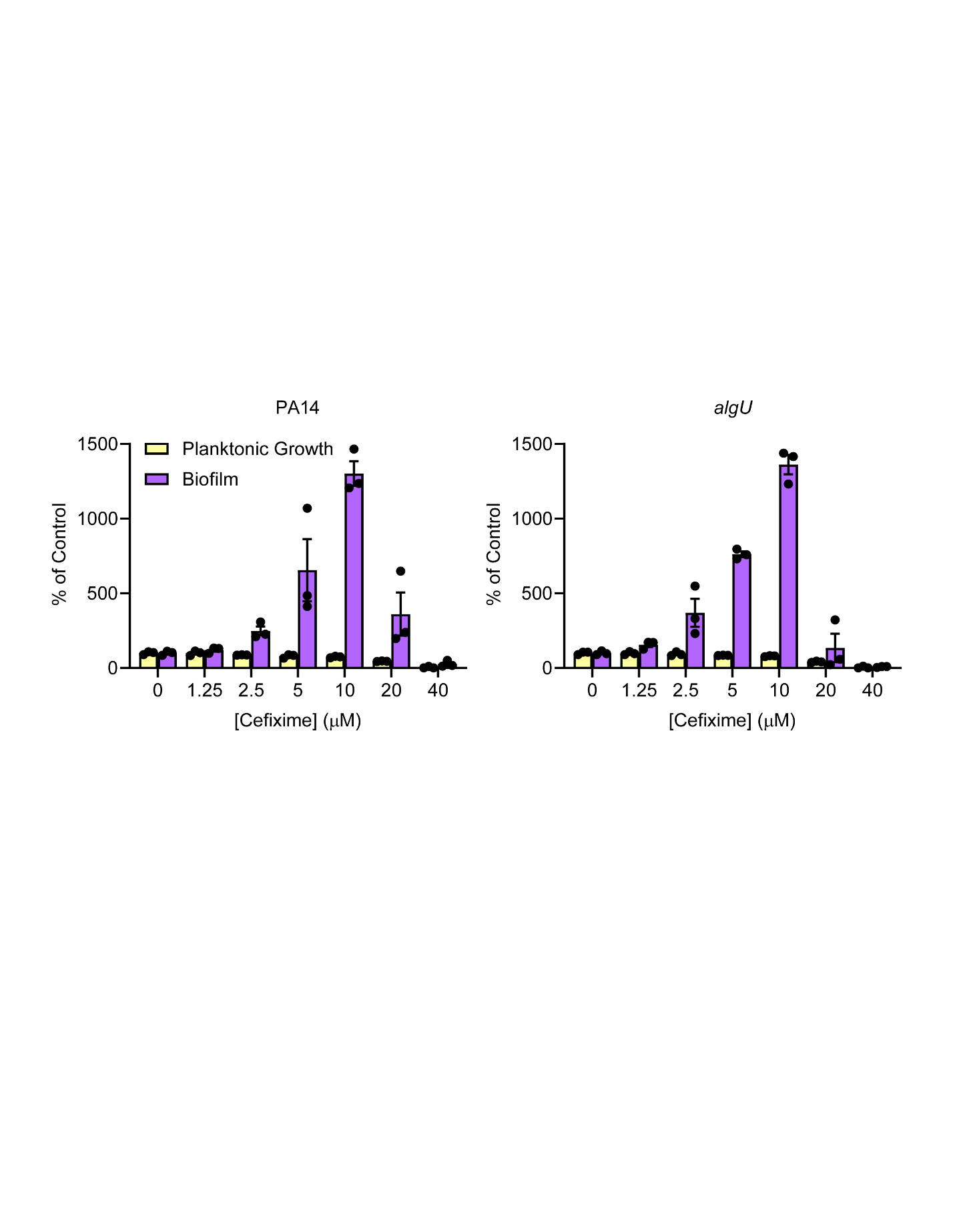

**Figure S2: Loss of *algU* does not impact biofilm stimulation by sub-MIC cefixime.** Treatment with sub-MIC cefixime stimulates biofilm formation in *P. aeruginosa* PA14 (left) and an isogenic *algU* deletion mutant (right). Planktonic growth (OD_600_, yellow) and biofilm (A_600_, purple) are reported as percentage of the untreated treated control. Two biological replicates were performed with 3 technical replicates for each, and a representative biological replicate is shown with the circle or triangle symbols representing individual data points. Error bars represent the standard error of the mean.
